# Global Quantitative Mapping of Enhancers in Rice Genome by STARR-seq

**DOI:** 10.1101/465716

**Authors:** Jialei Sun, Na He, Longjian Niu, Yingzhang Huang, Wei Shen, Yuedong Zhang, Li Li, Chunhui Hou

**Affiliations:** Department of Biology, Southern University of Science and Technology, Shenzhen 518055, China; Department of Bioinformatics, Huazhong Agriculture University, Wuhan 430070, China; Hubei Key Laboratory of Agricultural Bioinformatics, Huazhong Agriculture University, Wuhan 430070, China; Department of Biology, Nankai University, Tianjin 300071, China

**Keywords:** Rice; Enhancer, STARR-seq, Quantification, Functional analysis

## Abstract

Identification of enhancers has been a challenge in plants. STARR-seq measures enhancer activity of millions fragments in parallel. Here we present a global map of rice enhancers quantitatively determined using STARR-seq. Most enhancers are mapped within genes, especially at the 5’ untranslated regions (5’ UTR) and the coding sequences. Silent and low expressed genes in genomic regions enriched with transposable elements (TEs) are frequently found containing proximal enhancers. Analysis of enhancer epigenetic features at their endogenous loci revealed that most enhancers do not co-localize with DNase I hypersensitive sites (DHSs) and are lack of H3K4me1. Clustering enhancers by their epigenetic modifications revealed that about 40% of identified enhancers carry one or more epigenetic marks. Repressive H3K27me3 is frequently enriched with positive marks of H3K4m3 and/or H3K27ac, which together may bookmark poised enhancers. Intergenic enhancers were predicted based on the location of DHS relative to genes, which overlap poorly with functionally identified enhancers. In summary, enhancers were quantitatively identified by functional analysis in a model plant genome, which provides a valuable resource for further mechanistic studies in different biological contexts.

## Introduction

Gene expression is tightly regulated, which is critical for plant development and responses to changes in environment and hormones [1]. Promoters at transcription start site (TSS) are frequently considered sufficient for the initiation and elongation of transcription, but the level of promoter-driven expression is generally low [1]. High level of gene expression requires the participation of enhancers to increase the efficiency of transcriptional initiation and elongation to produce more mRNAs, though the exact mechanisms are still poorly understood.

Enhancers are generally believed to regulate their target genes through the recruitment of transcription factors (TFs) and function in a distance- and orientation-independent manner [2]. In mammalian genomes, one gene can be regulated simultaneously by multiple enhancers or by different enhancers in different tissues or at different developmental stages. Moreover, one enhancer can also regulate multiple genes [2-5]. The lack of specific location relationship with target genes makes the identification of enhancers more difficult than promoters.

The advancements in molecular biology and computational techniques have enabled the characterization of enhancers genome-widely based on epigenetic marks [6-15] or direct measurement of transcription enhancing activity of candidate sequences [16-22]. Intergenic enhancers have been predicted in *Arabidopsis* according to the location of DNase I hypersensitive site relative to proximal genes [23]. However, the arbitrary exclusion of DNase I sites within 1.5kb upstream of TSS and gene body may exclude substantial amount of potential enhancers. So far, only a handful of enhancers have been identified in plants [24-28], let alone a genome-wide annotation of enhancers based on functional analysis.

Measuring enhancer activity of millions fragments had been successfully achieved using STARR-seq in the *Drosophila melanogaster* and mammalian genomes [16, 18-20, 22]. We used STARR-seq to map potential enhancers globally in a rice genome. The accuracy of large scale quantification analysis of candidate sequences was demonstrated in a plant system. Furthermore, we revealed some epigenetic characteristics which may be unique for identified enhancers in rice genome. Finally, our results provided a regulatory element resource for further functional and mechanistic studies in different biological contexts.

## Results

### Global quantitative enhancer discovery using STARR-seq

To comprehensively identify sequences with enhancer activity, we constructed a reporter library from randomly fragmented genomic DNA of rice cultivar Nipponbare (*Oryza sativa L. ssp japonica*). The plasmids DNA of reporter library were transfected into protoplasts isolated from the stem of rice seedlings grown for 2 weeks in replicates and plasmids and mRNA were isolated 16 hours later. Sequencing libraries of plasmid and mRNA were generated and sequenced on an Illumina platform (**Figure 1A**). For two transfections, 15.5 million and 30.6 million (sTable 1) independent fragments were recovered with a median length of ~670 base pairs from input plasmid libraries (sFigure 1A) by Illumina paired-end sequencing. It covered ~90% of both repetitive and non-repetitive genome sequences with at least one unique fragment (sFigure 1C and 1E). For the cDNA libraries generated from isolated mRNAs, 6.1 million and 13.7 million of independent fragments were produced (sTable 1). The libraries were quality checked and the enrichment of cDNA over input plasmid was determined for each 600 bin and potential enhancers were identified (sFigure 2-6) and an example was shown in **Figure 1B**.

**Figure 1.**
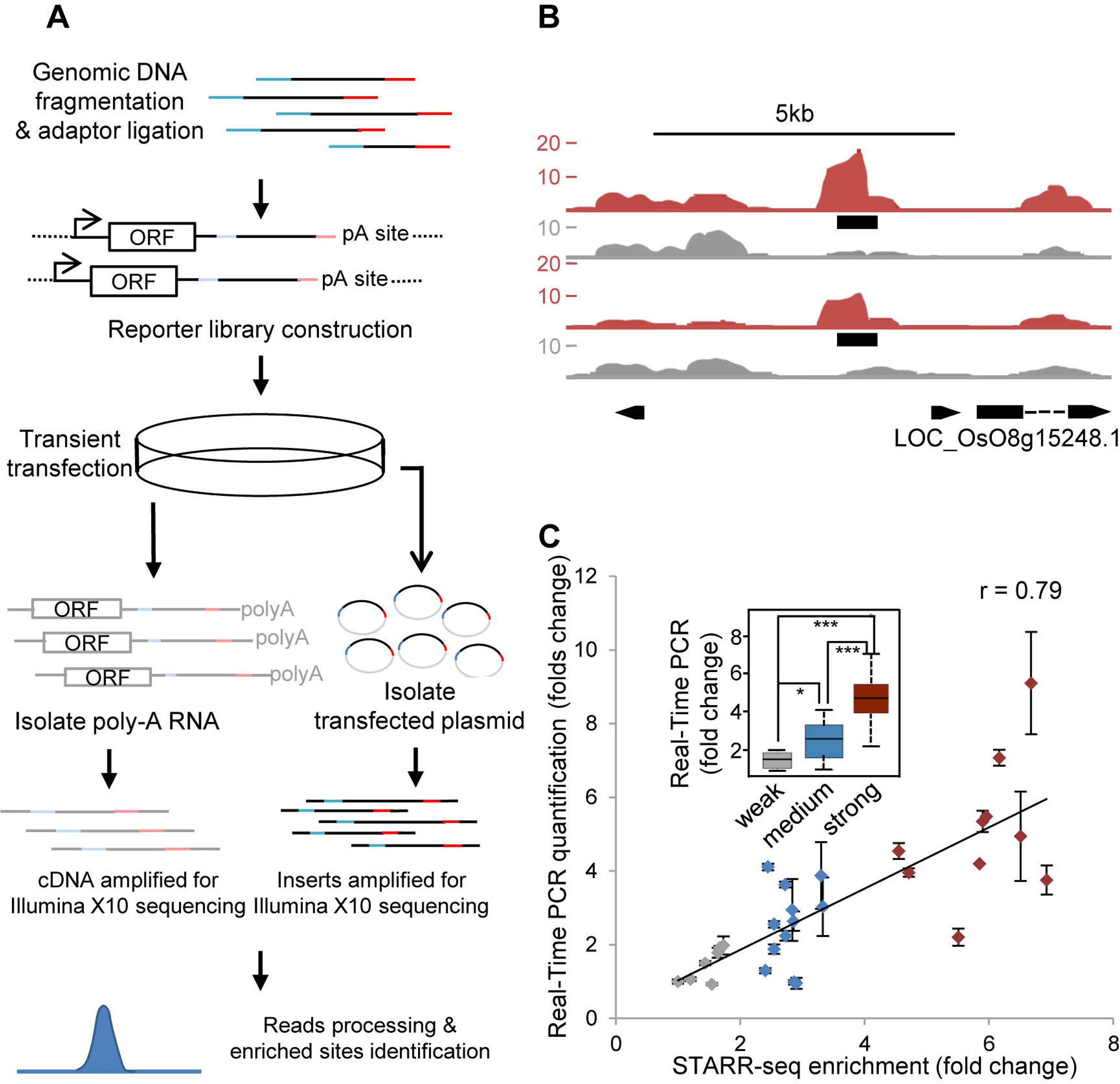
Genome-wide quantitative enhancer discovery. **A**. STARR-seq work flow. CMV 35s mini promoter was used to drive basal expression of reporter gene [ORF, open-reading frame; pA site, polyadenylation site]. **B**. STARR-seq cDNA (red) and input plasmid (grey) fragment densities in a genomic locus. Black boxes denote identified enhancers in two replicates. **C**. STARR-seq and Real-time PCR quantification are linearly correlated: r, Pearson correlation coefficient; Error bars indicate two independent biological replicates; Inset, the same data depicted in box plot; *p<0.05, ***p<0.001, Wilcoxon rank-sum test.

This method yielded 15,208 and 12,210 regions (**Figure 2A**) that were significantly enriched [p<0.001] from two replicates, respectively. Pearson correlation coefficient for the two replicates is 0.604 (sFigure 4), which indicates that STARR-seq in plant system is reproducible. Only enriched peaks identified in both biological replicates, totally 9,642 sites (sList 1), were kept for further analysis (**Figure 2A**). To validate the identified peaks, we tested 22 sites (sList 2) chosen across a wide range of enrichments by cloning them into luciferase reporter vector and quantified the reporter gene expression by reverse transcription and real-time PCR (RT-PCR) and normalized against the expression of co-transfected Renilla reporter gene. STARR-seq enrichment and RT-PCR quantification were strongly related from the entire activity range for enhancers from weak to strong (Pearson correlation coefficient, r = 0.79) (**Figure 1C**). The activity of the low, medium and strong enhancers also showed significant difference (p<0.05 and p<0.001 for comparisons between different groups, Wilcoxon test) (**Figure 1C**).

**Figure 2.**
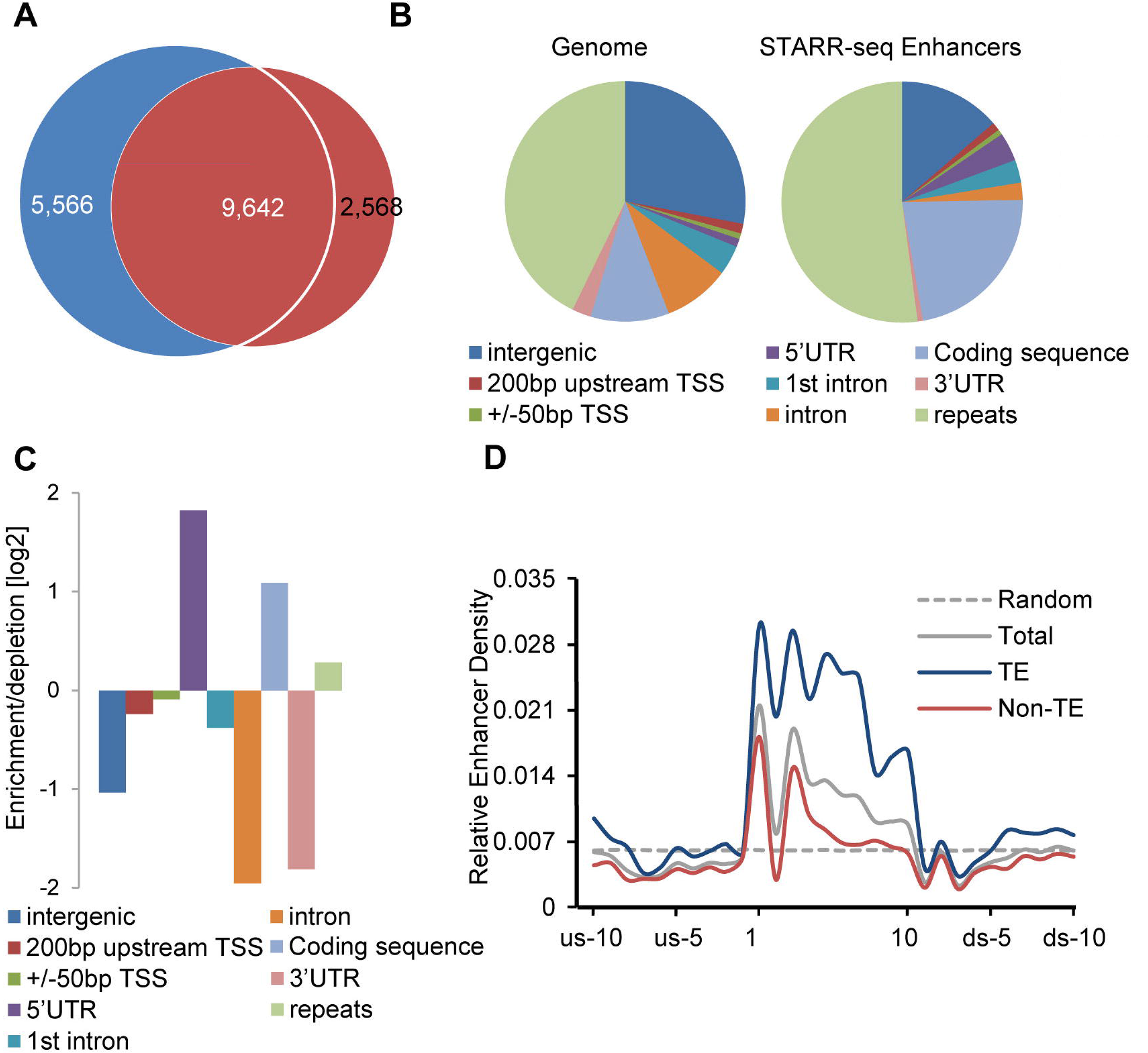
Genomic distribution of STARR-seq identified enhancers. **A.** Enhancers identified by two independent STARR-seq experiments. 9,642 enhancers were discovered in both replicates. **B** and **C**. Distribution and relative enrichment of identified enhancers in the rice genome. **D**. Distribution of enhancers relative to the gene body. TSS (transcription start site) and TTS (transcription termination site) are aligned and gene bodies are divided into 10 equal bins. Extended from TSS to upstream and from TTS to downstream are 10 bins of equal size (240bp, determined by dividing the median gene size by 10), respectively.

### Enhancers are enriched within gene body

STARR-seq identified enhancers in *Drosophila* genome are mostly located within genes and at promoter regions, especially in the introns (55.6%). Only 22.6% are in intergenic regions. *Drosophila* enhancers are significantly underrepresented in repetitive sequences [16]. Different from *Drosophila*, a little more than 50% of enhancers are mapped to repetitive sequences related to TEs (**Figure 2B**) and nearly all of these enhancers (4,831/5,020) contain repetitive elements including SINE, LINE, LTR, DNA transposones, satellite DNA and simple repeats (sTable 2). Different from animal genome, repetitive sequences in rice genome are gene-rich. The enrichment of enhancers in TE-related sequences in rice may be consistent with theories that TEs may regulate gene expression or even give rise to new genes during evolution [29, 30]. Identified enhancers are overrepresented in the 5’ untranslated regions (5’UTR) and coding sequences, while obviously underrepresented in introns (**Figure 2C**), which are strikingly different from the distribution pattern of enhancers in the *Drosophila* genome [16]. We divided the rice genome in two type, TE (repetitive sequences enriched with TEs) and non-TE (non-repetitive sequences) regions and analyzed the enhancer distribution relative to genes in these two types of sequences. Overall, the enhancer distribution patterns are quite similar in both TE and non-TE regions independent of the difference in gene activity in these two types of sequences (**Figure 2D**). Enhancers locate mostly within genes at the 5’ prime and gradually drop to background level toward the 3’ end of gene (**Figure 2D**).

### Proximal enhancers are absent from most genes in the rice genome

Majority enhancers are mapped within or close to genes, which suggests that proximal regulation by enhancer may be a prevalent choice in the rice genome. Compared to the total number of annotated and predicted genes (~56,000) [31], the number of enhancers is relatively low (9,642), less than 0.2 enhancer per gene averagely, suggesting most genes may be not directly regulated by enhancers in proximity (within TSS-5kb to TTS+5kb). Further analysis shows that 28.6% of genes (15,997) (**Figure 3A**) with at least one enhancer in proximity, suggesting one enhancer may be in close distance to multiple genes.

**Figure 3.**
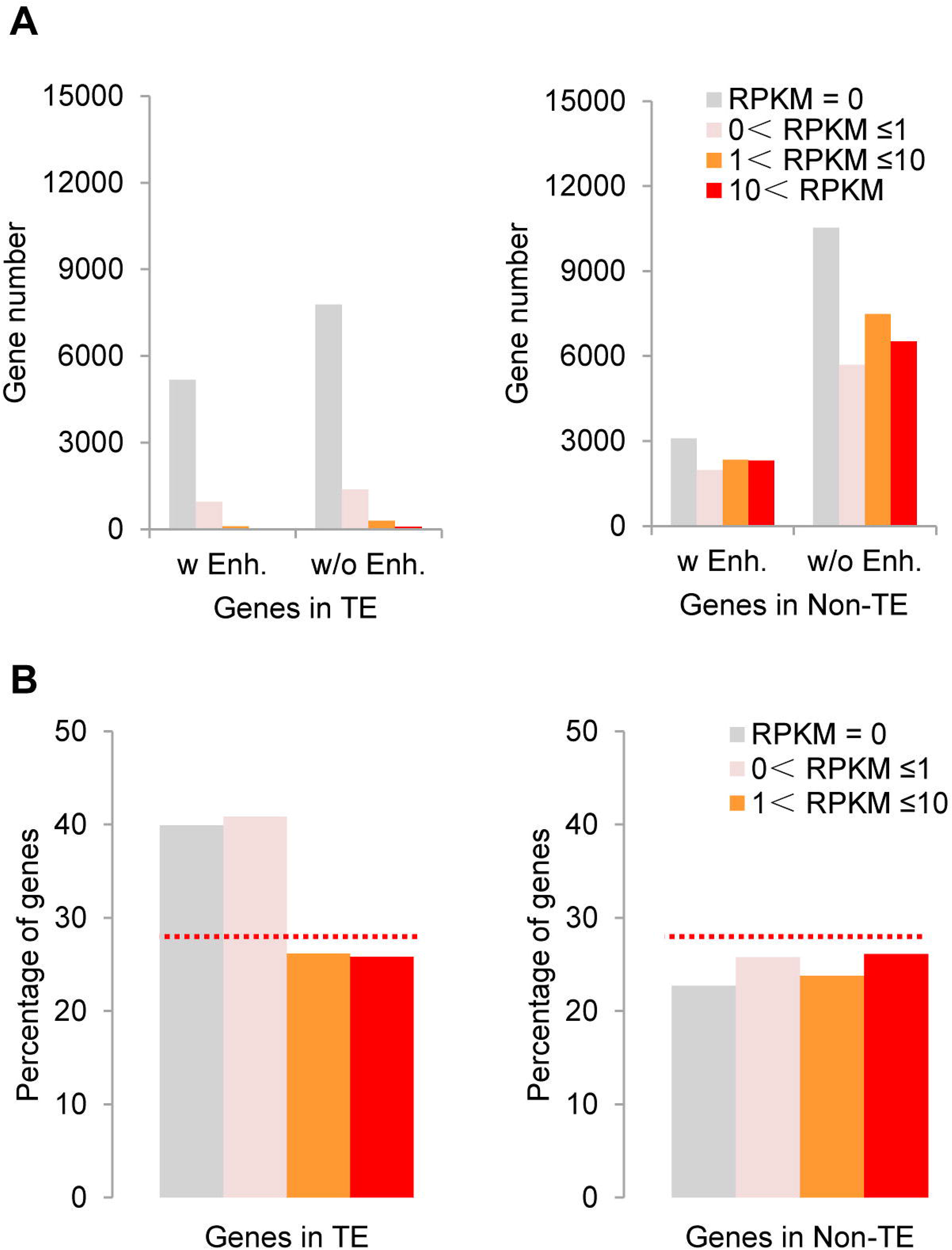
Presence of identified enhancers in proximity to genes of different expression level. **A**. Number of genes expressed at different levels with or without enhancers in proximity in TE (left) or non-TE regions (right), respectively. Enhancers within the range from TSS-5kb to TTS+5kb are defined as in proximity to genes. **B**. Percentage of genes expressed at different levels in TE (left) or non-TE regions (right) with or without enhancers in proximity. Silent, RPKM=0; Low, 0﹤RPKM≦1; Medium, 1﹤ RPKM≦10; High, RPKM﹥10.

We further separated genes into four groups according to their expression level (silent, low, medium and high) and calculated the percentage of genes with enhancer in proximity for each group. The percentage of genes with enhancer in proximity changes little for genes of different expression level in non-TE regions, and is slightly increased for genes in silence or expressed at low level in TE regions (**Figure 3B**). These results suggest STARR-seq enhancers at their endogenous loci are not enriched at actively transcribed genes, consistent with the episomal nature of the reporter plasmids.

### Genes in repetitive sequences are enriched with enhancers

Repetitive sequences related to TEs are generally transcriptionally inactive, which make up 42.8% of the rice genome (**Figure 2B**) and harbors 15,839 (28.3%) of total genes (**Figure 3A**, left panel). However, 52.1% of identified enhancers are located in TE regions (**Figure 2B**). These together suggest that genes in repetitive sequences are more enriched with enhancers. Actually, 39.6% of genes (6,277) located in TE regions contain at least one enhancer in proximity (**Figure 3A**, left panel). 96.7% of genes (15,318) in TE regions are of low expression level or silent and 40.1% of them (6,141) contain at least one enhancer in proximity. For genes of higher transcription levels or genes in non-TE regions, the percentage of genes with proximal enhancer is lower than the average for total genes (**Figure 3B**, right panel). Gene ontology (GO) analysis showed that genes in TE regions with enhancers in proximity occupy the top 10 most significantly enriched categories, which are related to DNA replication, integration, nucleotide binding, etc. (sFigure 7).

### Most identified enhancers are not DNase I hypersensitive sites (DHSs)

DNase I hypersensitivity has been generally believed to be associated with sequences of activities, one possibility is transcription enhancing. 8.8% of all identified enhancers overlap with DHSs (**Figure 4A**). Genes with proximal enhancer overlapping with DHS in both TE and non-TE regions show higher expression level (**Figure 4B**). Similarly, only about 13.6% of enhancers identified in the human genome co-localize with DHSs (sTable 3) [20]. Differently, 48.5% of enhancers identified in *Drosophila* co-localize with DHSs (sTable 3) [16]. The lack of DHS for the majority of enhancers proximal to actively transcribed genes may suggest that hypersensitivity to DNase I digestion may not be a universal mark for all functional enhancers in rice genome.

### H3K4me1 is not a ubiquitous enhancer mark in rice genome

Histone modification mark H3K4me1 has been frequently used for enhancer prediction, which is enriched at enhancers identified by STARR-seq in both *Drosophila* and human genomes [16, 20]. However, for enhancers identified in rice, H3K4me1 is even less enriched than DHS, only ~330 sites show strong signal of H3K4me1 (**Figure 5A**). H3K4me1 is nearly completely absent from predicted enhancers based on DHS locations in the genome (**Figure 6A** and sFigure 8).

### H3K4me3 is enriched at many identified enhancers

H3K4me3 is generally associated with actively transcribed genes. A significant portion of enhancers locate inside active genes at 5’ UTR and in coding sequences (**Figure 2D** and **Figure 3**). Consistently, H3K4me3 is found obviously enriched for enhancers identified, more obvious than any other histone mark or DHS (**Figure 4C & 4D**, **Figure 5A** and sFigure 8).

### H3K27ac is an enriched enhancer mark

Another histone modification mark frequently used for the prediction of active enhancer is H3K27ac, which has also been shown significantly enriched at enhancers identified by STARR-seq in both *Drosophila* and human genomes [16, 20]. Different from H3K4me1, H3K27ac is also enriched for identified enhancers in rice genome, the signal is especially strong for enhancers associated with DHS or located within non-TEs (red lines in **Figure 4C & 4D**, **Figure 5A** and sFigure 8). Even enhancers not associated with DHS show slightly enrichment of H3K27ac (blue line in **Figure 4C**).

**Figure 4.**
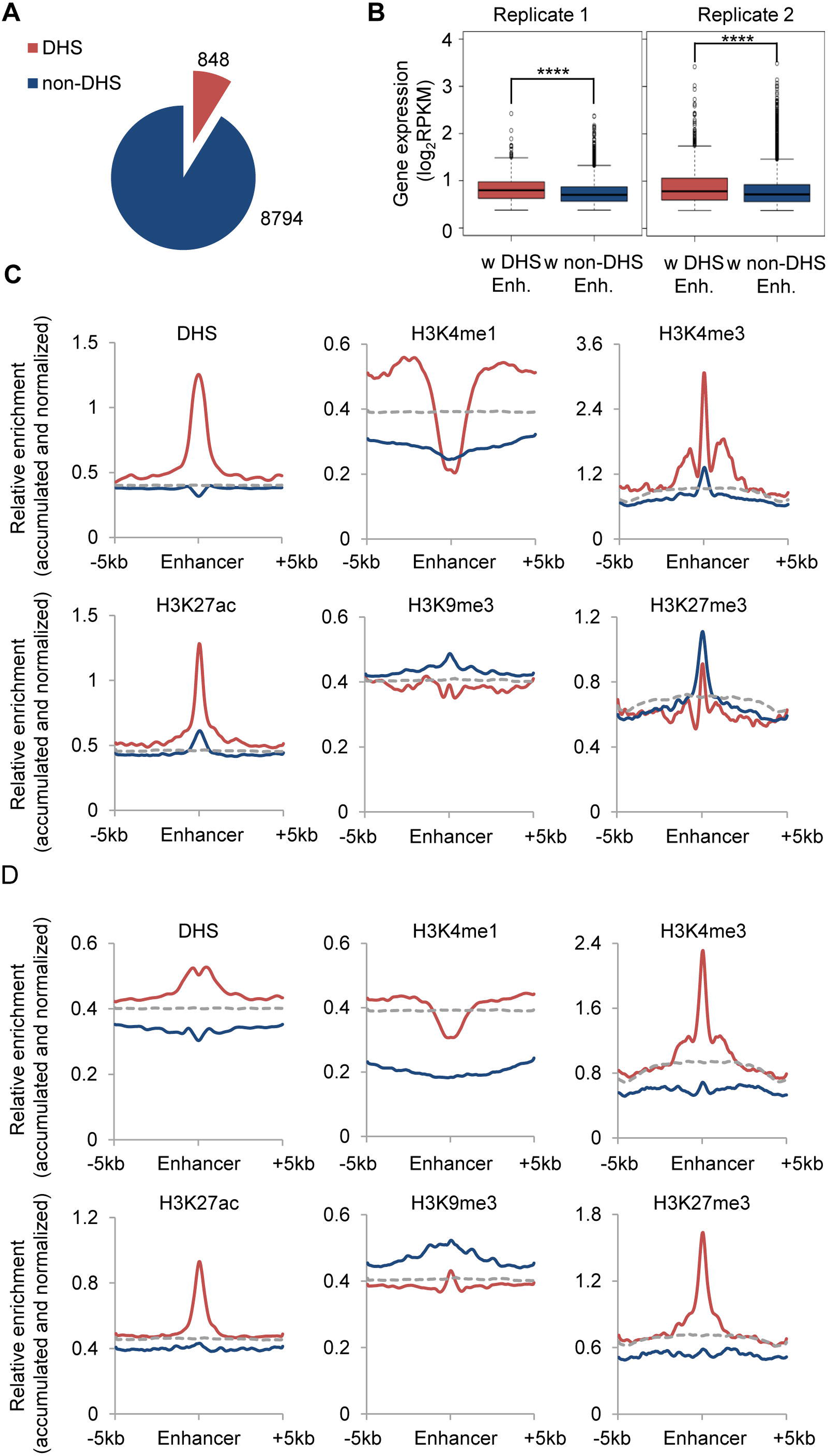
Epigenetic characteristics of identified enhancers at their endogenous loci. **A**. Numbers of identified enhancers overlap with DNase I hypersensitive sites, DHSs. **B**. Expression of genes in TE or non-TE regions with proximal enhancers overlapping with DHS or not; ****p<10^−10^, Wilcoxon rank-sum test. **C**. Epigenetic signal enrichment for enhancers overlapping with DHSs (red) or not (blue). **D**. Epigenetic signal enrichment for enhancers in non-TE (red) and TE (blue) regions, respectively.

### H3K27me3 is enriched at a significant number of identified enhancers

H3K27me3 is mostly associated with repressed genes. Surprisingly, we found that a significant number of identified enhancers are enriched with H3K27me3 (**Figure 4C & 4D**), which is generally absent from enhancers identified in *Drosophila* and human genomes [16, 20]. Most of these enhancers are enriched strongly with H3K4me3 and H3K27ac as well (**Figure 5A** and sFigure 8). That whether these enhancers are poised or actively regulating genes is difficult to determine. Different from H3K27me3, H3K9me3 is nearly completely absent from identified enhancers (**Figure 5A** and sFigure 8).

**Figure 5.**
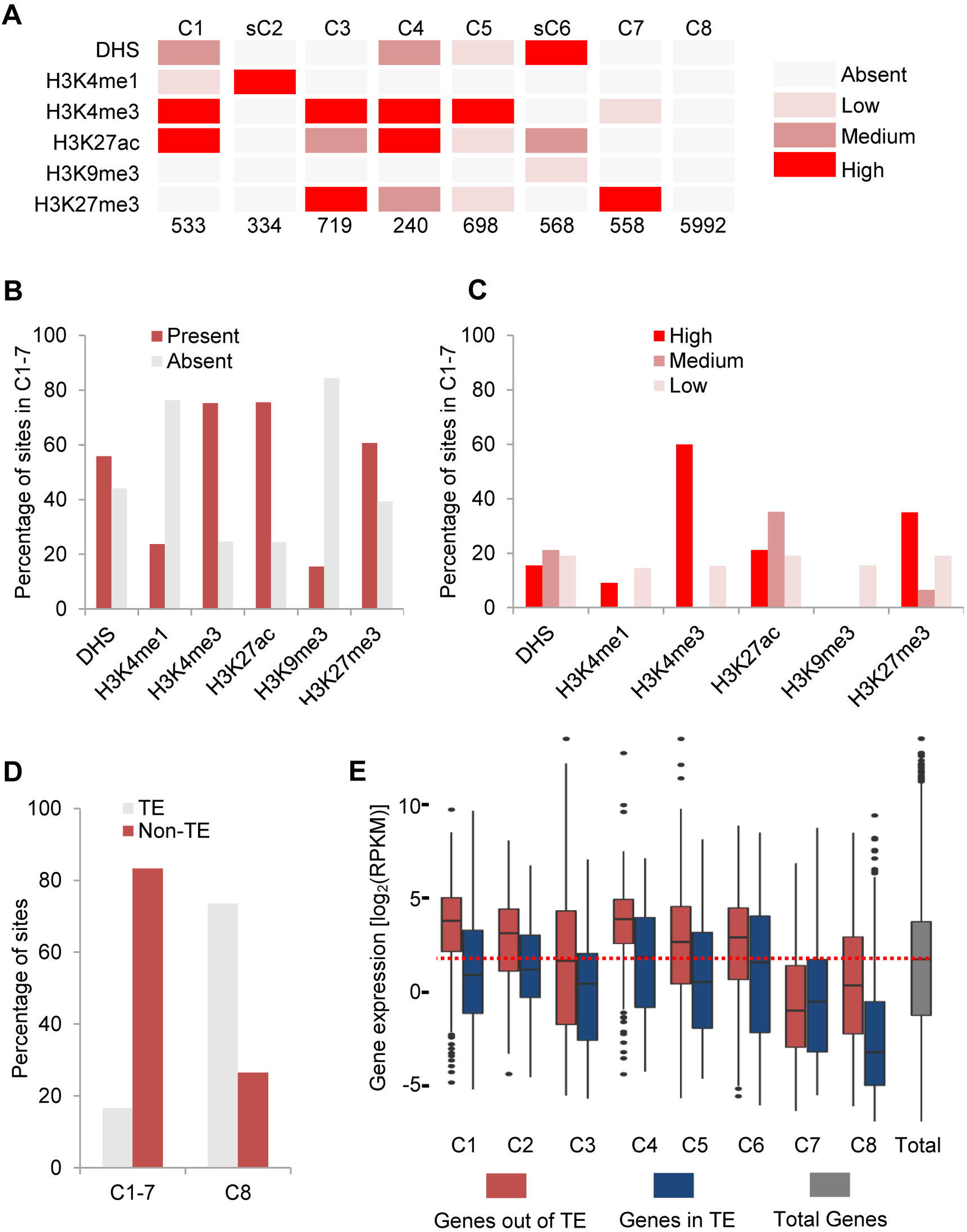
Epigenetically clustered STARR-seq enhancers. **A**. Identified enhancers are grouped into 8 clusters based on the signal strength of 6 epigenetic marks. Numbers of enhancers in each cluster are shown at the bottom. Overall signal density was ranked: absent (close to background level) as light grey box, low as light red, medium as dark red and high as bright red. **B**. Percentage of identified enhancers in each cluster with (red) or without (light grey) indicated epigenetic mark. **C**. Identified enhancers with indicated epigenetic mark in each cluster were further divided into three sub-classes based on the average signal density, percentage of enhancers of low, medium and high signal density are shown. **D**. Percentage of cluster 1-7 and C8 in TE (light grey) or non-TE (red) regions. **E**. Expression of genes in TE (blue) or non-TE (red) regions with enhancers of different cluster in proximity. Genes in non-TE regions with enhancers of nearly all clusters (except C7) show significant (p<0.01) higher expression level than in TEs.

### Epigenetic clustering of identified enhancers

We further grouped all enhancers into eight clusters based on the signal strength of multiple epigenetic marks including DHS, H3K4me1, H3K4me3, H3K27ac, H3K9me3 and H3K27me3 (**Figure 5A** and sFigure 8). Enhancers of cluster 1-7 (C1, sC2, C3-5, sC6 and C7; total 3,650 enhancers, 37.9% of total enhancers identified; sC2 and sC6 are unique for STARR-seq identified enhancers) are enriched with at least one epigenetic mark (**Figure 5A** and sFigure 8). However, 62.1% of enhancers (5,992) are devoid of any analyzed epigenetic mark (C8, data not shown). H3K4me3 is strongly enriched for 4 clusters of enhancers which at the same time are enriched with different levels of H3K27ac (**Figure 5A** and sFigure 8). Overall, H3K4me3 and H3K27ac are mostly enriched, DHS and H3K27me3 are medially enriched, H3K4me1 and H3K9me3 are least enriched for enhancers of C1-7 (**Figure 5B & 5C** and sFigure 8). About 17% and 83% of C1-7 enhancers locate inside TE and non-TE regions, respectively (**Figure 5D**). Accordingly, majority of C8 enhancers (73%) are associated with TE regions (**Figure 5D**). Genes associated with each cluster of enhancers are expressed at significantly (p<0.01, Wilcoxon test) higher level in non-TE regions than in TE regions except for cluster 7 (p=0.916) (**Figure 5E**). Enhancers in cluster 7 are enriched with H3K27me3 (**Figure 5A**).

### Predicted enhancers based on DHS location overlap little with STARR-seq identified enhancers

Enhancers had been predicted based on chromatin accessibility in *Arabidopsis* [23]. We followed the published methods and defined a DHS as enhancer if it locates beyond 1.5kb upstream of TSS and at the same time is not in a gene body. By this method, 13,770 out of total 37,168 DHSs were predicted as enhancers (**Figure 6A,** sList 3). Only 20% of them are in TE regions and 80% of them are in non-TE regions (**Figure 6B**), consistent with the fact that the repetitive sequences are mostly in closed chromatin states with low accessibility. Due to the different distribution pattern, DHS predicted enhancers overlap with only a few STARR-seq enhancers (**Figure 6C**). Strikingly, clustering of predicted enhancers (DHS signal omitted and the C8 is different from STARR-seq enhancers C8, dC2 and cC6 are also unique) showed that even less of them are enriched with histone modification marks (**Figure 6D**). Overall, the expression of genes with proximal DHS predicted enhancer in TE and non-TE regions differ little (p>0.05, Wilcoxon test) except those being associated with enhancers in clusters of dC2 and dC6 (p<0.01, Wilcoxon test, **Figure 6E** and sFigure 8). Moreover, except dC6 and C8, genes with enhancers in proximity are not expressed significantly higher than the middle level of total genes (**Figure 6E**).

**Figure 6.**
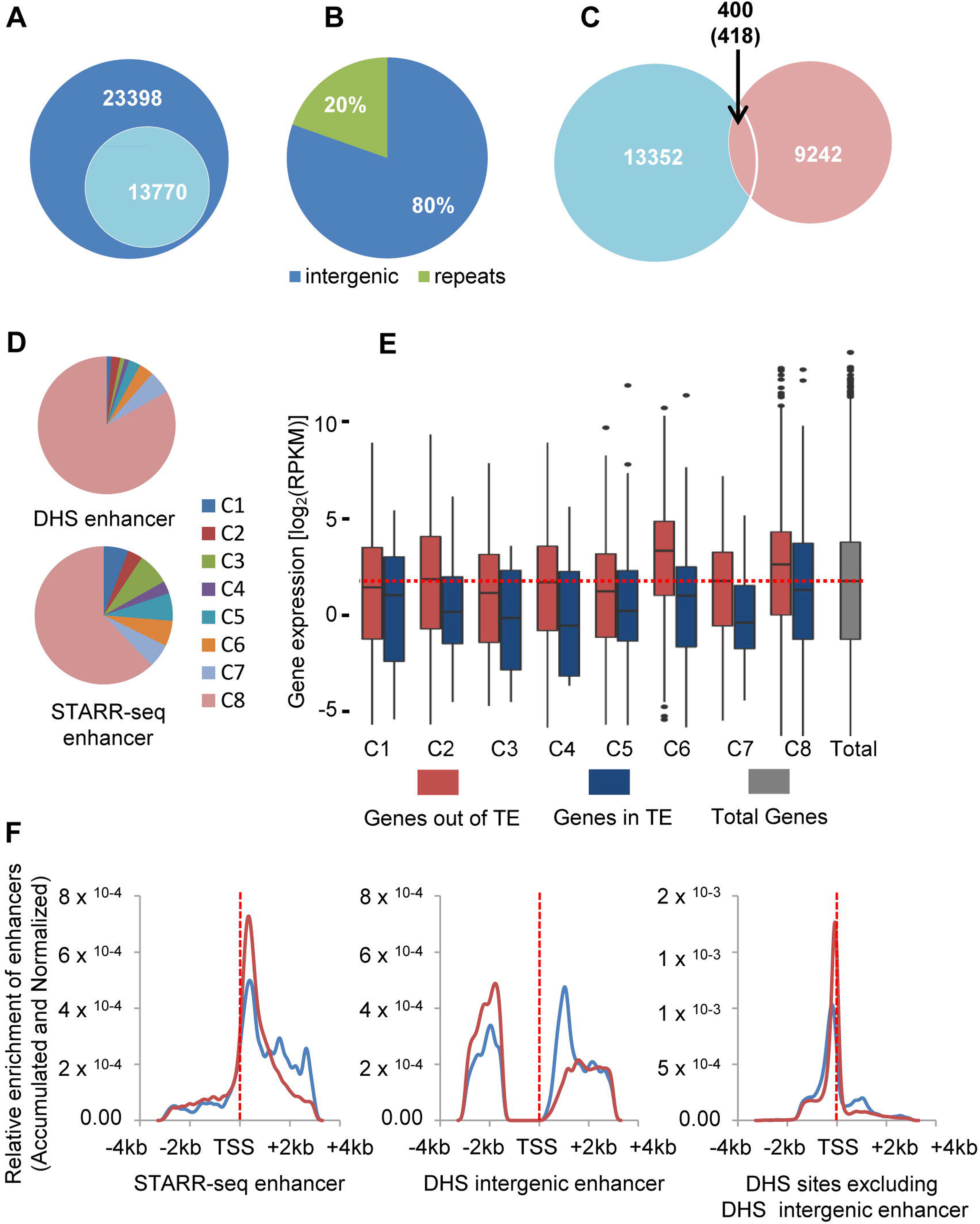
Predicted enhancers based on DHSs location to genes. **A**. Numbers of predicted enhancers (13,770) according to DHSs location and of DHSs with predicted enhancer excluded (23,398). **B**. Distribution of predicted enhancers in the genome. **C**. Overlap of predicted enhancers and STARR-seq enhancers. **D**. Pie charts of clustered enhancers. For predicted enhancers, DHS signal was omitted for clustering. **E**. Expression of genes in TE (blue) or non-TE (red) regions with enhancers of each cluster in proximity. Genes in non-TE regions with enhancers of clusters 2, 6, 7 and 8 show significant (p<0.01) higher expression than genes in TE regions. **F**. Relative enrichment of identified enhancers, predicted enhancers and DHS sites excluding predicted enhancers around the TSS. Red and blue lines show sites in non-TE or TE regions, respectively.

We further compared the distribution of STARR-seq enhancers, predicted DHS enhancers and other DHS sites (with DHS predicted enhancers excluded, non-predicted enhancer DHSs, sList 4). These three groups of elements show sharply different distribution patterns relative to the TSS of genes (**Figure 6F**). As previously shown, STARR-seq enhancers are mostly enriched within gene body favoring the 5’ (**Figure 6F**, left). Most DHSs are closely positioned to TSSs. In fact, 91% of all DHSs in rice genome are mapped within TSS+/-5kb regions, and similarly, 80.9% in *Drosophila* genome (sFigure 9). DHS predicted enhancers (13,770 out of total 37,168 DHSs genome-wide, 37%) are at least 1.5kb upstream away from TSS and out of gene body (**Figure 6F**, middle). For non-predicted enhancer DHSs (23,398 out of total 37,168, 63%) (sFigure 10a), they are predominantly enriched at upstream of TSS where promoters are located (**Figure 6F**, right), and only 1,347 of them overlap with STARR-seq enhancers (sFigure 10b). Due to close positioning to TSSs, significant portion of these DHSs overlap with sequences within the 200 bp upstream of TSS, +/-50bp of TSS and the 5’ UTR regions (sFigure 10c), which is dramatically different from both DHS predicted and STARR-seq enhancers, respectively. Moreover, clustering on epigenetic modifications revealed that 50.5% of these sites are enriched with at least one type of modifications and most of them are enriched with H3K4me3, an active mark for active genes (sFigure 8, right). Though the majority of DHS sites are located not far from genes, these sites are less enriched inside gene body, where most of the identified enhancers are located.

## Discussion

Enhancer prediction relies heavily on chromatin epigenetic mark. Though several histone modifications (H3K4me1 for most enhancers, H3K27ac for active enhancers, and coexistence of H3K27me3 and H3K4me1 for poised enhancers) have been frequently used for enhancer analysis, this method may fail to predict enhancers in chromatin devoid of histone modifications. Moreover, enhancers predicted by epigenetic marks are difficult to verify by genetic methods at large scale. To measure enhancer activity of candidate sequences independent from chromatin context, STARR-seq was invented [16] and had been successfully applied to the enhancer analysis in both the *Drosophila* and human genomes [16, 20]. However, a genome-wide functional analysis of enhancers for a plant genome has not been reported, which severely limits our understanding of the nature and functional mechanisms of plant enhancers.

Previous work predicting enhancers by chromatin sensitivity to DNase I digestion excluded arbitrarily DHSs within 1.5kb upstream of TSS and in gene body [23]. In *Drosophila*, the majority of STARR-seq enhancers are actually located within gene body or in proximity to genes [16]. Similarly, our STARR-seq analysis of rice genome also shows that the majority of enhancers are localized within the gene body. The consistent observation of enhancer enrichment in the gene body in two evolutionarily far-separated genomes may suggest that most genes could be regulated by DNA elements built in their sequences (**Figure 7**). Furthermore, it will be interesting to see if enhancers in genes can also activate other genes separated by long distance. A capture Hi-C using identified enhancers as anchors may reveal if in plant genome genes separated far away can be co-regulated by enhancers located within the gene body.

**Figure 7.**
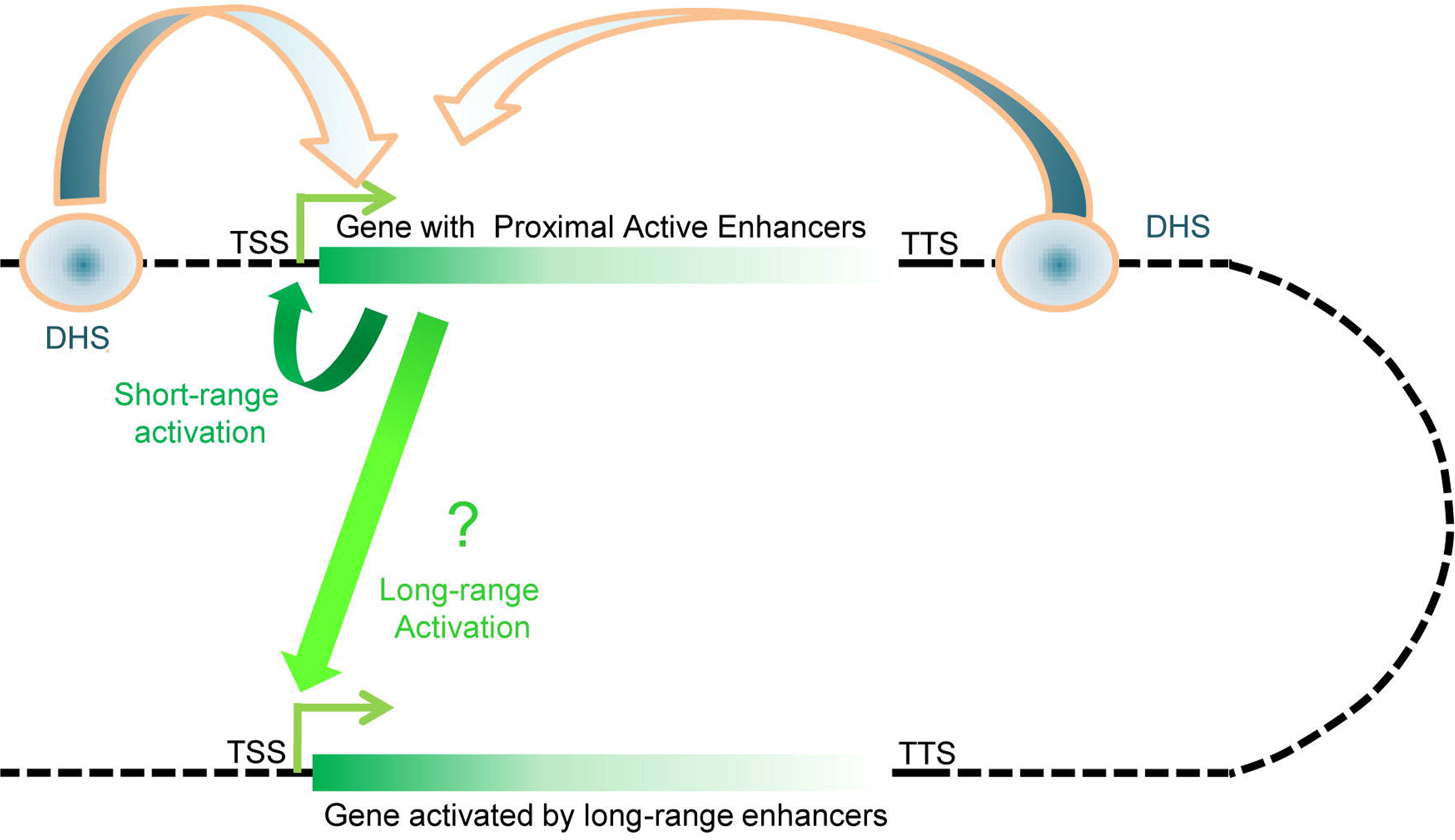
Proposed enhancer function model in rice genome. Gene body is shown in green. Density of identified enhancers gradually decreases within gene body from 5’ TSS to the 3’ TTS. Genes can be self-regulated by enhancers located within. Enhancers in genes may also activate other genes separated far away. Predicted enhancers are mostly located away from genes and if they activate genes or not requires systematic experimental validation.

Our analysis also reveals several unexpected features of identified enhancers in the rice genome. First, the majority of STARR-seq enhancers do not overlap with DHSs. DNase I hypersensitivity can be associated to any open and relaxed chromatin region, including insulators and other protein binding sites. Predicting enhancers relies solely on DHS location relative to the gene may fail to exclude DHSs which may possess non-enhancer activity.

Second, though H3K4me1 has been used to predict enhancers in mammals, it is actually devoid from most STARR-seq enhancers and DHSs (independent of their location) in rice genome. The fact that H3K4me1 is depleted from most DHSs further confirms that H3K4me1 may not be the ideal mark of enhancers in rice genome.

Third, many identified enhancers are enriched with H3K27me3. The majority of these enhancers are co-enriched with active chromatin marks of H3K4me3 and H3K27ac. Co-existence of H3K27me3 and H3K4me3 can be explained by the theory that many enhancers are poised. Surprisingly, H3K27ac and H3K27me3 co-exist at quite a few enhancer sites. These two modifications are supposed to be mutually exclusive. At this time, it is difficult to rule out possibilities that histones at these enhancers may be modified differently in different cells, or even differently on different allele in the same cell. In either case, it requires further careful analysis to reveal the underlying causes of this intriguing observation.

In summary, we presented a comprehensive enhancer activity map generated by quantitative measurement using STARR-seq for an important model plant. This method is especially suitable for functional analysis of small genomes. Successful characterization of enhancers in different cell types will help to improve our understanding of the nature of enhancers in the plant genome and sheds new lights into the elusive functional mechanisms of enhancers at large.

## Materials and methods

### Screening vector

For STARR-seq in rice cells, we constructed a screening vector based on the plasmid pPBI221 backbone and modified by introducing several sequences, which include a CMV 35S mini promoter, an intron and a GFP sequence, which are arranged sequentially and are underlined in the following DNA sequence.

ggcttggggttagtctcaccagtcacagaaagcatcttacggatggcatgacagtaagagaattatgcagtgctgccataaccatgagtgataacactgcggccaacttacttctgacaacgatcggaggaccgaaggagctaaccgcttttttgcacaacatgggggatcatgtaactcgccttgatcgttgggaaccggagctgaatgaagccataccaaacgacgagcgtgacaccacgatgcctgtagcaatggcaacaacgttgcgcaaactattaactggcgaactacttactctagcttcccggcaacaattaatagactggatggaggcggataaagttgcaggaccacttctgcgctcggcccttccggctggctggtttattgctgataaatctggagccggtgagcgtgggtctcgcggtatcattgcagcactggggccagatggtaagccctcccgtatcgtagttatctacacgacggggagtcaggcaactatggatgaacgaaatagacagatcgctgagataggtgcctcactgattaagcattggtaactgtcagaccaagtttactcatatatactttagattgatttaaaacttcatttttaatttaaaaggatctaggtgaagatcctttttgataatctcatgaccaaaatcccttaacgtgagttttcgttccactgagcgtcagaccccgtagaaaagatcaaaggatcttcttgagatcctttttttctgcgcgtaatctgctgcttgcaaacaaaaaaaccaccgctaccagcggtggtttgtttgccggatcaagagctaccaactctttttccgaaggtaactggcttcagcagagcgcagataccaaatactgttcttctagtgtagccgtagttaggccaccacttcaagaactctgtagcaccgcctacatacctcgctctgctaatcctgttaccagtggctgctgccagtggcgataagtcgtgtcttaccgggttggactcaagacgatagttaccggataaggcgcagcggtcgggctgaacggggggttcgtgcacacagcccagcttggagcgaacgacctacaccgaactgagatacctacagcgtgagctatgagaaagcgccacgcttcccgaagggagaaaggcggacaggtatccggtaagcggcagggtcggaacaggagagcgcacgagggagcttccagggggaaacgcctggtatctttatagtcctgtcgggtttcgccacctctgacttgagcgtcgatttttgtgatgctcgtcaggggggcggagcctatggaaaaacgccagcaacgcggcctttttacggttcctggccttttgctggccttttgctcacatgttctttcctgcgttatcccctgattctgtggataaccgtattaccgcctttgagtgagctgataccgctcgccgcagccgaacgaccgagcgcagcgagtcagtgagcgaggaagcggaagagcgcccaatacgcaaaccgcctctccccgcgcgttggccgattcattaatgcagctggcacgacaggtttcccgactggaaagcgggcagtgagcgcaacgcaattaatgtgagttagctcactcattaggcaccccaggctttacactttatgcttccggctcgtatgttgtgtggaattgtgagcggataacaa tttcacacaggaaacagctatgaccatgattacgccaagcttgggccgctcgagccgcccgggcga**<u>cttcgcaagacccttcctctatataaggaagttcatttcatttggagag</u>**tccggatccattaacctgcaggtaaatttctagagctctggtagaaatctgagg**<u>gtaaatttctagtttttctccttcattttcttggttaggacccttttctctttttatttttttgagctttgatct ttctttaaactgatctattttttaattgattggttatggtgtaaatattacatagctttaactgataatctgattactttat ttcgtgtgtctatgatgatgatgatagttacag</u>**aaccgacgactcgtc**<u>atgacatccactttgcctttctctccacagg tgtccactcccaggtccaactgcaggtcgcctgcaggcttaaccatggctagcaaaggagaagaactcttcactggagttgtcccaattcttgttgaattagatggtgatgtaaacggccacaagttctctgtcagtggagagggtgaaggtgatgcaacatacggaaaacttaccctgaagttcatctgcactactggcaaactgcctgttccctggccaacactagtcactactctgtgctatggtgttcaatgcttttcaagatacccggatcatatgaaacggcatgactttttcaagagtg ccatgcccgaaggttatgtccaggaaaggaccatcttcttcaaagatgacggcaactacaagacacgtgctgaagtcaagtttgaaggtgatacccttgttaatagaatcgagttaaaaggtattgacttcaaggaagatggcaacattctgggacacaaattggaatacaactataactcacacaatgtatacatcatggcagacaaacaaaagaatggaatcaaagcgaacttcaagacccgccacaacattgaagatggaagcgttcaactagcagaccattatcaacaaaatactccaattggcgatggccctgtccttttaccagacaaccattacctgtccacacaatctgccctttcgaaagatcccaacgaaaagagagaccacatggtccttcttgagtttgtaacagctgctgggattacacatggcatggatgaactgtataa ctga</u>**tctagcgcatgcaccgggccctggtagaaatctgaggaaccgacgactcgtctgtacaacactctttccctacacg acacacgtctgaactccagtctgtaattcacgcgtccagggatcgttcaaacatttggcaataaagtttcttaagattgaatcctgttgccggtcttgcgatgattatcatataatttctgttgaattacgttaagcatgtaataattaacatgtaatgcatgacgttatttatgagatgggtttttatgattagagtcccgcaattatacatttaatacgcgatagaaaacaaaatatagcgcgcaaactaggataaattatcgcgcgcggtgtcatctatgttactagatcgggccagtcgacccaagatcttccccggaattcactggccgtcgttttacaacgtcgtgactgggaaaaccctggcgttacccaacttaatcgccttgcagcacatccccctttcgccagctggcgtaatagcgaagaggcccgcaccgatcgcccttcccaacagttgcgcagcctgaatggcgaatggcgcctgatgcgtctagagtattttctccttacgcatctgtgcggtatttcacaccgcatatggtgcactctcagtacaatctgctctgatgccgcatagttaagccagccccgacacccgccaacacccgctgacgcgccctgacgggcttgtctgctcccggcatccgcttacagacaagctgtgaccgtctccgggagctgcatgtgtcagaggttttcaccgtcatcaccgaaacgcgcgagacgaaagggcctcgtgatacgcctatttttataggttaatgtcatgataataatggtttcttagacgtcaggtggcacttttcggggaaatgtgcgcggaacccctatttgtttatttttctaaatacattcaaatatgtatccgctcatgagacaataaccctgataaatgcttcaataatattgaaaaaggaagagtatgagtattcaacatttccgtgtcgcccttattcccttttttgcggcattttgccttcctgtttttgctcacccagaaacgctggtgaaagtaaaagatgctgaagatcagttgggtgcacgagtgggttacatcgaactggatctcaacagcggtaagatccttgagagttttcgccccgaagaacgttttccaatgatgagcacttttaaagttctgctatgtggcgcggtattatcccgtattgacgccgggcaagagcaactcggtcgccgcatacactattctcagaatgacttggttgagtactcaccagtcacagaaaagcatcttacggatggcatgacagtaagagaattatgcagtgctgccataaccatgagtgataacactgcggccaacttacttctgacaacgatcggaggaccgaaagagctaaccgcttttttgcacaacatgg

### Generation of input (screening) libraries

We extracted genomic DNA from two weeks rice seedlings. About 125 µg of genomic DNA was diluted to 50ng/µl and fragmented by sonication (Scientz II-D). DNA fragments (500bp-800bp length) were size-selected, end repaired, 5’-phosphorylated and 3’ dA-tailed with VAHTS Mate Pair Library Prep Kit for Illumina^®^ (VAHTS; cat. no. ND104). VAHTS Adapters for Illumina was ligated to about 6µg of DNA fragments using VAHTS Mate Pair Library Prep Kit for Illumina^®^ (VAHTS; cat. no. ND104) following the manufacturer’s protocol. Adaptor ligated DNA was purified by using GeneJET PCR Purification Kit (Thermo Scientific; cat. no. K0702) then was amplified by TransStart FastPfu Fly DNA Polymerase (Transgen; cat. no. AP231) with Illumina sequencing primers (fw: ACACTCTTTCCCTACACGACG & rev: GACTGGAGTTCAGACGTGTGC).

We obtained linear pPBI221 by PCR amplification (95°C for 5min; then 25 cycles of 95°C for 20s, 57°C for 20s and 72°C for 4min; Forward primer: ACACGTCTGAACTCCAGTCTGTAATTC & Reverse primer: GTCGTGTAGGGAAAGAGTGTTGTACA). Circular plasmids was removed by digestion with DpnI (NEB; cat. no. R0176) at 37°C for 30min. PCR products were recovered with E.Z.N.A.^®^ Gel Extraction Kit (Omega; cat. no. D2500). The adaptor ligated genomic DNA were recombined to linearized screening vector with ClonExprress II One Step Cloning Kit (Vazyme; cat. no. C112). For each reaction, 100ng linearized vector and 40ng adaptor ligated DNA was used in a total volume of 20µl. 70 reactions were performed in total. Each of 140 tubes (100µl each) of Trans1-T1 Phage Resistant Chemically Competent Cell (Transgen; cat. no. CD501) was transformed with 10µl of DNA, according to the manufacturer’s protocol. 140 transformation reactions were pooled, transferred to 4L LB_AMP_ medium, and grown to OD 0.8. Plasmids were purified using E.Z.N.A.^®^ Endo-Free Plasmid Maxi Kit (Omega; cat. no. D6926) then quantified.

### Protoplast preparation

The rice cultivar Nipponbare (*Oryza sativa L. ssp japonica*) seeds were moistened with water in Petri dishes at room temperature to promote germination (3days) and then were transferred to culture box at 37°C for about 24h, and then transplanted into soil. The rice plants were grown at 28°C with a relative humidity around 60%. The light/dark photoperiod is 12/12 h by using a bank of 400 W metal halide lamps. The light intensity is 400 µmol m^−2^s^−1^ at the plant height. We collected 2-week-old rice seedlings for the all experiments.

Protoplasts isolation is described below. Stem tissues from 80-120 rice seedlings were used. 5 seedlings were cut together into approximately 0.5 mm strips using sharp razors in 0.6M D-mannitol on Petri dishes. The strips were then digested (1.5% Cellulase RS, 0.6 M mannitol, 0.1% BSA, 0.75% Macerozyme R-10, 3.4mM CaCl2, 10 mM MES, pH 5.7), and then vacuumed at −50kPa in bottle for 30 min. After that, the bottle was put on oscillator for 4-5h in dark with gentle shaking (45rpm) at 28°C. After digestion, protoplasts were filtered through 300 micron nylon meshes and washed 3-5 times with W5 solution (125 mM CaCl2, 154 mM NaCl, 5 mM KCl, 2 mM MES, pH 5.7). Protoplasts were pelleted by centrifugation at 80g for 5min. After another two wash with W5 solution (equal volume), protoplasts were then resuspended in MMG solution (15 mM MgCl2, 0.6M mannitol, 4 mM MES, pH 5.7), and placed on ice for 30 min. Protoplasts were centrifuged at 80g again for 5 min and finally resuspended at a concentration of 1 × 10^7^ cells per ml using the MMG solution, determined by using a hematocytometer. All procedures were carried out at room temperature.

### Plasmid library transformation

For each sample, 30-40μg of plasmid was mixed with 100μL protoplasts (~1× 10^6^ cells) in a 2 ml tube with 110μl of freshly prepared PEG solution [40% (W/V), 0.1 M CaCl_2_, 0.6 M mannitol] added. Gently mix, then incubate in dark at 28°C for 15 min. Add 800μl of W5 solution slowly. Invert gently to mix well, then centrifugate immediately at 80g for 5 min. The pelleted protoplasts were resuspended gently in 800μl of W5 solution. Finally, protoplasts transfected were cultured at 28°C for 16 h in dark. Reporter GFP signals were checked under fluorescent microscopy.

### Reporter cDNA libraries construction for Illumina sequencing

16hrs post-transfection, cells were washed in W5 solution twice and concentrated. Total RNA was extracted using TransZolTM Up Plus RNA Kit (ER501). polyA+ RNA fraction was isolated using VAHTS mRNA Capture Beads (N401). 5U DNase I (NEB M0303S) was used to digest DNA at 37°C for 20 min.

Synthesis of first strand cDNA was carried out with TransScript One-Step gDNA Removal and cDNA synthesis SuperMix (AU311 50°C for 30 min and 85°C for 5s) using a specific primer (GACTGGAGTTCAGACGTGTGC) and 8ul of RNA in 20 reactions. All reactions were pooled. The total amount of reporter cDNA was amplified for Illumina sequencing by a 2-step nested PCR strategy using the TransStart^®^ FastPfu Fly DNA Polymerase (AP231-12).

In the first PCR (98°C, 2min; then 10-12 cycles of 98 °C for 30s, 58°C for 30s, 72°C for 30s; and then 72°C for 5min), 10-30ng cDNA were used for amplification with 2 reporter-specific primers (Fw: 5’GACTGGAGTTCAGACGTGTGC3’ & Rev: 5’GAGCTCTGGTAGAAATCTGAGGGTGTC3’), one of which spans the RNA splice junction of GFP intron. PCR products were purified by GeneJET PCR Purification Kit (K0701) and eluted in 20-30ul Elution Buffer (EB). Purified PCR product as template for the second PCR (98°C for 2min; followed by 6-10 cycles of 98°C for 30s, 65°C for 30s, 72°C for 30s; and then 72 °C for 5min) with TransGen^®^ Biotech FastPfu Fly DNA Polymerase (AP231) and with VAHTS^TM^ DNA Adapters for Illumina^®^ (N302). PCR products were purified by GeneJET PCR Purification and eluted in 20-30 ul EB.

### Transfected plasmid library construction for Illumina sequencing

When the PolyA RNA was capture, keep the total RNA (without mRNA) supernatant, add 10ul RNase A TransGen^®^ Biotech GE 101 20mg/ml at 37°C for 30-60min, and the plasmid was purified by GeneJET PCR Purification Kit and eluted in 50 ul EB. Purified plasmid as template for the second PCR (98°C for 2min; followed by 8-12 cycles of 98°C for 30s, 65°C for 30s, 72°C for 30s; and then 72 °C for 5min) with TransGen^®^ Biotech FastPfu Fly DNA Polymerase and with VAHTS™ DNA Adapters for Illumina^®^ (N302). PCR products were purified by GeneJET PCR Purification and eluted in 20-30 ul EB. All the libraries were sequenced on Illumine X Ten platform.

### Computational analysis

#### Mapping Illumina sequences

For 150mer paired-end reads generated by Illumina X Ten platform, bowtie2 [32] was used to map them to the Nipponbare reference genome (IRGSP1.0). The default parameters were used except parameter “–no-discordant –X 2000”. After mapping, we used the samtools [33] to filter the mapped reads and kept only uniquely mapped reads with parameter “view -bS -f 2 -q 5”.

#### Computational processing of STARR-seq data

Unique fragments were used to compute the genome coverage in non-repeat region and the enrichment of cDNA signal over plasmid input. Detailed procedures to identify enhancers were described in Arnold et al (2013) [16]. And the program used in this experiment was an R package named BasicSTARRseq. We performed bonferroni correction to correct p values. Peaks are kept with p-value smaller than 0.001 and enrichment fold signal over 1.3. Only enhancers found in both replicates were kept and used for further analysis. The overlapped enhancers were merged by bedtools merge [34].

#### SATRR-seq enhancers overlap with DNase I hypersensitive sites (DHSs)

We consider it is an overlap if the middle point of an enhancer falls in a DHS site. By this definition, 848 enhancers were found overlapping with DHS. DHS data were generated by Zhang et al. [35]. DHS open sites were called by MACS1.4 [36] with default parameter.

#### DHS predicted enhancers according to their location relative to genes

Enhancers were predicted in the rice genome following the definition published previously for enhancer prediction in Arabidopsis [23]. DHSs located in intergenic regions (>1.5 kb upstream of TSS and downstream of transcription terminal site, TTS) were arbitrarily defined as enhancers. Total 37,168 DHSs were called by MACS1.4 in the whole rice genome. We used bedtools intersect to filter the DHSs in intergenic regions and 13,770 DHSs were considered as predicted enhancers.

#### Correlation analysis between different libraries

Reads inside consecutive 500bp bins were used to calculate the correlation between different libraries. Bins with value of zero were excluded from the calculation. We used multiBamSummary and plotCorrelation in deeptools [37] with parameter: –binSize 500 –ignoreDuplicates, and -corMethod pearson –skipZeros. Pearson correlation coefficients were calculated.

#### Coverage analysis

Only unique fragment data were used to compute the read coverage at each position for both repetitive and non-repetitive genome sequences. We used bedtools to calculate the coverage ratio.

#### Enhancer distribution in the genome

We used MSU 7.0 to annotate the rice genome in the following categories: repeat, coding sequence, core promoter (+/- 50 bp TSS around the TSS), 5’ UTR, 3’ UTR, 1st intron, intron, 200bp upstream TSS, intergenic. Each enhancer was sorted into different category if its summit falls within sequence of a specific category. The distance between enhancer and proximal TSS was computed by bedtools closest command.

#### Analysis of GC-content effect of plasmid/cDNA libraries

We calculated the GC content for each non-overlapping 100bp bins in the non-repeat genome. Sequences were divided into 100 bins from 20% to 80% according to their GC content. The cDNA/plasmid mean read coverage in each bin was also calculated. The softwares used are makewindows, nuc, coverage of bedtools suit.

#### Intersecting genomic coordinates

We used bedtools suite of programs to intersect genomic coordinates for STARR-seq enhancers, DHS enhancers and the closest TSS to a specific enhancer.

#### STARR-seq, DHS-seq, and chromatin mark enrichment

Histone modification data, including H3K4me3, H3K27me3 were retrieved from GSE19602 [38], and the H3K27ac, H3K9me3 were retrieved from GSE79033 [39], the H3K4me1 was retrieved from GSA under accession no. PRJCA000387 [40]. DHS data were from GSE26734 [35]. All the histone modifications were mapped to the reference genome IRGSP 1.0 using bowtie2 with parameter –no-discordant -X 2000 for paired-end data and –no-discordant for single-end data, and only the uniquely mapped reads were kept with duplicates removed by picardtools. The histone mark peaks were called by MACS1.4 with default parameters. Adapter was removed from the DHS sequencing data by cutadapt before mapping. Duplicates were removed with picardtools, and peaks were called by MACS1.4. Reads density at each position was normalized for comparison. 10,000 randomly selected regions of 700bp were used as control, and repeated for at least 10 times to calculate the mean value. The enrichment of chromatin mark and DHS were showed in a 5kb window with the enhancer peak positioned in the center. STARR-seq enhancer, DHS enhancer and DHSs except predicted enhancers were divided into two groups depending on if they were located in repeat regions. STARR-seq enhancers were also divided into two groups according to their location relative to DHS. We used the R package EnrichedHeatmap [41] to plot the histone modifications (H3K4me1, H3k27ac, H3k4me3, H3K9me3, K3K27me3), and DHS enrichment for all groups with the center of analyzed elements positioned at middle point and extended to upstream and downstream by 5kb, respectively.

#### Determine the proximity of enhancer to genes

An enhancer is defined in proximity of a gene if it is located within the range of gene body +/- 5kb region. Genes were separated in two groups, TE and non-TE according to their location in either TE or non-TE regions. The number of TE gene is 15,839, and the number of Non-TE gene is 39,961. The tools we used is BedTools. Gene expression data are from Zhang et al. (2016) [31].

#### GO analysis of genes with enhancer in proximity

We used agriGO V2.0 [42] for the GO enrichment analysis.

#### Histone modification at enhancers

Relation between histone modifications, DHS and enhancers were analyzed. 3,650 out of total 9,642 STARR-seq enhancers overlap with at least one type of histone mark or DHS. 2,331 out of 13,770 DHS enhancers overlap with at least one kind of histone mark, while 11,819 out of 23,398 DHS peaks (with DHS enhancers excluded) overlap with at least one type of histone mark.

#### Enhancer categorizations based on epigenetic marks

Enhancers were classified into 8 clusters based on histone marks, DHS and relative enrichment of these marks. K-Means in Cluster3.0 [43] was used. Expression level of proximal genes with enhancers of each category was analyzed.

#### Distribution of STARR-seq enhancers relative to the gene body

TSSs and TTSs were aligned and sequences between were equally divided into 10 bins. Another 2.4kb was extended upwards and downwards from TSS and TTS, respectively. The 2.4kb extended regions were divided into 10 equal sized bins as well. The size of 2.4kb was arbitrarily chosen for it is the median size of all genes in rice genome. 55,000 regions of 2.4kb were randomly generate by bedtools random command with parameter -n 55000 -l 2400, which were repeated for 100 times. Genes were separated in TE and non-TE groups. We counted the enhancer number in each bin and normalized against the total number of genes in each category or 55,000 for random control.

#### Percentage of genes with enhancers in proximity

Genes were divided into four groups based on their expression level. High, RPKM >10; Medium, 1＜RPKM≦10; Low, 0＜RPKM≦1 and Silent. Genes were separated into two groups, TE and non-TE, to compare.

#### Motif analysis

For each group of elements, we submitted their sequences to the MEME-ChIP web server19 for de novo motif finding, using default parameters. The database we used was JASPAR CORE (2018) 20 plant database. Identified motifs are enclosed in supplementary List 5.

#### Statistical analysis

We used R for all statistical analysis.

#### Data Availability

All data are available under the accession number GSE121231.

## Authors’ contributions

JS constructed the reporter library and validated identified sites. LN designed and monitored the experiments. YZ participated in cell preparation, transfection and sequencing library preparation. NH carried out bioinformatics analysis. WS processed the raw data and participated in bioinformatics analysis. YZ helped with reporter library construction. LL advised on data analysis. CH designed, monitored the experiments and wrote the manuscript with input from all authors.

## Acknowledgements

We gratefully acknowledge financial support from the National Natural Science Foundation of China (31571347 to CH and 31771430 to LL), Guangdong Science and Technology Department (2016A030313642 to CH), Shenzhen Science and Technology Innovation Commission (JCYJ20150529152146478 to CH), Huazhong Agricultural University Scientific & Technological Self-innovation Foundation (to LL) and the Young Thousand Talent Program (to CH). We thank Dr. Shengtao Hou for manuscript editing.

## Supplementary materials

Figures S1 to S10

Tables S1 to S3

Lists S1 to S5

**Supplementary Figure 1 Plasmid and cDNA libraries**

**A** and **B**. Fragment sizes of STARR-seq input (top left) and cDNA libraries (top right), the median fragment sizes are indicated. **C** and **D**. Coverage of the rice non-repetitive genome by independent fragments (cumulative), **C** for plasmid library, **D** for cDNA library. **E** and **F**. Coverage of the rice repetitive genome by independent fragments (cumulative), **E** for plasmid library, **F** for cDNA library. More than 90% of the genome is covered by more than one fragment in input plasmid library.

**Supplementary Figure 2 GC content analysis for plasmid and cDNA libraries**

The genome sequence was binned according to GC-content (x axis). Each boxplot shows the read depth of all positions within the respective windows, revealing that STARR-seq input shows similar pattern as the cDNA library.

**Supplementary Figure 3** Correlation analysis of all sequenced libraries. See Methods for analysis description.

**Supplementary Figure 4 Correlation analysis of enhancers from two replicates**

For all enhancers identified by two independent biological replicates, the correlation of their activity was reasonably high with a Pearson correlation coefficient r = 0.604.

**Supplementary Figure 5 Enhancer distribution relative to their strength**

**Supplementary Figure 6 Number of unique fragments supporting enhancer identification** Percentage of enhancers versus the number of independent genomic fragments per enhancer.

**Supplementary Figure 7 GO analysis of genes mostly enriched**

Top ten categories of enriched genes with identified enhancer in proximity. Most of them are located within TE regions and are involved in the biological process of DNA synthesis.

**Supplementary Figure 8 Enhancer and DHSs clustering on epigenetic signals**

STARR-seq enhancers, DHS predicted enhancers and the remaining DHSs are clustered. Note that the two groups of DHSs were clustered without the values of DNase I hypersensitivity. sC2, sC6 are unique for STARR-seq enhancers. dC2, dC6 are unique for the two groups of DHS elements.

**Supplementary Figure 9 Distribution of DHSs relative to TSSs**

The relative DHSs enrichment in Drosophila and rice genomes are calculated in moving 500bp bins normalized against total number of bins at each position. 0 on X axis corresponding to TSS.

**Supplementary Figure 10 DHSs with predicted enhancer sites excluded**

A. Total DHSs are divided into two groups, 13,770 sites as predicted enhancers, 23,398 sites within TSS-1.5kb and in gene body. B. Overlap of STARR-seq enhancers and non-predicted enhancer DHSs sites. C. Relative enrichment of non-predicted enhancer DHSs sites in different categories of genome sequences.

**Supplementary Table 1 Sequencing statistics of libraries**

**Supplementary Table 2 STARR-seq enhancers in TE regions overlap with different types of repetitive sequences**

**Supplementary Table 3 Percentage of STARR-seq enhancers overlapping with DHS**

**Supplementary List 1 STARR-seq identified enhancers**

**Supplementary List 2 Real-Time PCR validated enhancer sites**

**Supplementary List 3 Predicted enhancers according to DHS location**

**Supplementary List 4 non-predicted-enhancer DHSs**

**Supplementary List 5 Identified motifs for different groups of elements**

## References

[1] Marand AP, Zhang T, Zhu B, Jiang J. Towards genome-wide prediction and characterization of enhancers in plants. Biochim Biophys Acta Gene Regul Mech. 2017;1860(1):131–9.

[2] Bulger M, Groudine M. Functional and mechanistic diversity of distal transcription enhancers. Cell. 2011;144(3):327–39.

[3] Kieffer-Kwon KR, Tang Z, Mathe E, Qian J, Sung MH, Li G, et al. Interactome maps of mouse gene regulatory domains reveal basic principles of transcriptional regulation. Cell. 2013;155(7):1507–20.

[4] Li G, Ruan X, Auerbach RK, Sandhu KS, Zheng M, Wang P, et al. Extensive promoter-centered chromatin interactions provide a topological basis for transcription regulation. Cell. 2012;148(1-2):84–98.

[5] Tang Z, Luo OJ, Li X, Zheng M, Zhu JJ, Szalaj P, et al. CTCF-Mediated Human 3D Genome Architecture Reveals Chromatin Topology for Transcription. Cell. 2015;163(7):1611–27.

[6] Whyte WA, Orlando DA, Hnisz D, Abraham BJ, Lin CY, Kagey MH, et al. Master transcription factors and mediator establish super-enhancers at key cell identity genes. Cell. 2013;153(2):307–19.

[7] Visel A, Taher L, Girgis H, May D, Golonzhka O, Hoch RV, et al. A high-resolution enhancer atlas of the developing telencephalon. Cell. 2013;152(4):895–908.

[8] Hnisz D, Abraham BJ, Lee TI, Lau A, Saint-Andre V, Sigova AA, et al. Super-enhancers in the control of cell identity and disease. Cell. 2013;155(4):934–47.

[9] Zentner GE, Tesar PJ, Scacheri PC. Epigenetic signatures distinguish multiple classes of enhancers with distinct cellular functions. Genome Res. 2011;21(8):1273–83.

[10] Kim TK, Hemberg M, Gray JM, Costa AM, Bear DM, Wu J, et al. Widespread transcription at neuronal activity-regulated enhancers. Nature. 2010;465(7295):182–7.

[11] Creyghton MP, Cheng AW, Welstead GG, Kooistra T, Carey BW, Steine EJ, et al. Histone H3K27ac separates active from poised enhancers and predicts developmental state. Proc Natl Acad Sci U S A. 2010;107(50):21931–6.

[12] Visel A, Blow MJ, Li Z, Zhang T, Akiyama JA, Holt A, et al. ChIP-seq accurately predicts tissue-specific activity of enhancers. Nature. 2009;457(7231):854–8.

[13] Heintzman ND, Hon GC, Hawkins RD, Kheradpour P, Stark A, Harp LF, et al. Histone modifications at human enhancers reflect global cell-type-specific gene expression. Nature. 2009;459(7243):108–12.

[14] Heintzman ND, Stuart RK, Hon G, Fu Y, Ching CW, Hawkins RD, et al. Distinct and predictive chromatin signatures of transcriptional promoters and enhancers in the human genome. Nat Genet. 2007;39(3):311–8.

[15] Visel A, Rubin EM, Pennacchio LA. Genomic views of distant-acting enhancers. Nature. 2009;461(7261):199–205.

[16] Arnold CD, Gerlach D, Stelzer C, Boryn LM, Rath M, Stark A. Genome-wide quantitative enhancer activity maps identified by STARR-seq. Science. 2013;339(6123):1074–7.

[17] Inoue F, Ahituv N. Decoding enhancers using massively parallel reporter assays. Genomics. 2015;106(3):159–64.

[18] Vanhille L, Griffon A, Maqbool MA, Zacarias-Cabeza J, Dao LT, Fernandez N, et al. High-throughput and quantitative assessment of enhancer activity in mammals by CapStarr-seq. Nat Commun. 2015;6:6905.

[19] Dao LTM, Galindo-Albarran AO, Castro-Mondragon JA, Andrieu-Soler C, Medina-Rivera A, Souaid C, et al. Genome-wide characterization of mammalian promoters with distal enhancer functions. Nat Genet. 2017;49(7):1073–81.

[20] Liu Y, Yu S, Dhiman VK, Brunetti T, Eckart H, White KP. Functional assessment of human enhancer activities using whole-genome STARR-sequencing. Genome Biol. 2017;18(1):219.

[21] Muerdter F, Boryn LM, Woodfin AR, Neumayr C, Rath M, Zabidi MA, et al. Resolving systematic errors in widely used enhancer activity assays in human cells. Nat Methods. 2018;15(2):141–9.

[22] Liu S, Liu Y, Zhang Q, Wu J, Liang J, Yu S, et al. Systematic identification of regulatory variants associated with cancer risk. Genome Biol. 2017;18(1):194.

[23] Zhu B, Zhang W, Zhang T, Liu B, Jiang J. Genome-Wide Prediction and Validation of Intergenic Enhancers in Arabidopsis Using Open Chromatin Signatures. Plant Cell. 2015;27(9):2415–26.

[24] Yang W, Jefferson RA, Huttner E, Moore JM, Gagliano WB, Grossniklaus U. An egg apparatus-specific enhancer of Arabidopsis, identified by enhancer detection. Plant Physiol. 2005;139(3):1421–32.

[25] Clark RM, Wagler TN, Quijada P, Doebley J. A distant upstream enhancer at the maize domestication gene tb1 has pleiotropic effects on plant and inflorescent architecture. Nat Genet. 2006;38(5):594–7.

[26] McGarry RC, Ayre BG. A DNA element between At4g28630 and At4g28640 confers companion-cell specific expression following the sink-to-source transition in mature minor vein phloem. Planta. 2008;228(5):839–49.

[27] Schauer SE, Schluter PM, Baskar R, Gheyselinck J, Bolanos A, Curtis MD, et al. Intronic regulatory elements determine the divergent expression patterns of AGAMOUS-LIKE6 subfamily members in Arabidopsis. Plant J. 2009;59(6):987–1000.

[28] Raatz B, Eicker A, Schmitz G, Fuss E, Muller D, Rossmann S, et al. Specific expression of LATERAL SUPPRESSOR is controlled by an evolutionarily conserved 3' enhancer. Plant J. 2011;68(3):400–12.

[29] Zhao D, Ferguson AA, Jiang N. What makes up plant genomes: The vanishing line between transposable elements and genes. Biochim Biophys Acta. 2016;1859(2):366–80.

[30] Hirsch CD, Springer NM. Transposable element influences on gene expression in plants. Biochim Biophys Acta Gene Regul Mech. 2017;1860(1):157–65.

[31] Zhang J, Luo W, Zhao Y, Xu Y, Song S, Chong K. Comparative metabolomic analysis reveals a reactive oxygen species-dominated dynamic model underlying chilling environment adaptation and tolerance in rice. New Phytol. 2016;211(4):1295–310.

[32] Langmead B, Salzberg SL. Fast gapped-read alignment with Bowtie 2. Nat Methods. 2012;9(4):357–9.

[33] Li H, Handsaker B, Wysoker A, Fennell T, Ruan J, Homer N, et al. The Sequence Alignment/Map format and SAMtools. Bioinformatics. 2009;25(16):2078–9.

[34] Quinlan AR, Hall IM. BEDTools: a flexible suite of utilities for comparing genomic features. Bioinformatics. 2010;26(6):841–2.

[35] Zhang W, Wu Y, Schnable JC, Zeng Z, Freeling M, Crawford GE, et al. High-resolution mapping of open chromatin in the rice genome. Genome Res. 2012;22(1):151–62.

[36] Zhang Y, Liu T, Meyer CA, Eeckhoute J, Johnson DS, Bernstein BE, et al. Model-based analysis of ChIP-Seq (MACS). Genome Biol. 2008;9(9):R137.

[37] Ramirez F, Ryan DP, Gruning B, Bhardwaj V, Kilpert F, Richter AS, et al. deepTools2: a next generation web server for deep-sequencing data analysis. Nucleic Acids Res. 2016;44(W1):W160–5.

[38] He G, Zhu X, Elling AA, Chen L, Wang X, Guo L, et al. Global epigenetic and transcriptional trends among two rice subspecies and their reciprocal hybrids. Plant Cell. 2010;22(1):17–33.

[39] Fang Y, Wang X, Wang L, Pan X, Xiao J, Wang XE, et al. Functional characterization of open chromatin in bidirectional promoters of rice. Sci Rep. 2016;6:32088.

[40] Pan X, Fang Y, Yang X, Zheng D, Chen L, Wang L, et al. Chromatin states responsible for the regulation of differentially expressed genes under (60)Co~gamma ray radiation in rice. BMC Genomics. 2017;18(1):778.

[41] Gu Z, Eils R, Schlesner M, Ishaque N. EnrichedHeatmap: an R/Bioconductor package for comprehensive visualization of genomic signal associations. BMC Genomics. 2018;19(1):234.

[42] Tian T, Liu Y, Yan H, You Q, Yi X, Du Z, et al. agriGO v2.0: a GO analysis toolkit for the agricultural community, 2017 update. Nucleic Acids Res. 2017;45(W1):W122–W9.

[43] de Hoon MJ, Imoto S, Nolan J, Miyano S. Open source clustering software. Bioinformatics. 2004;20(9):1453–4.

